# Electrical microstimulation evokes saccades in posterior parietal cortex of common marmosets

**DOI:** 10.1101/693580

**Authors:** Maryam Ghahremani, Kevin D. Johnston, Liya Ma, Lauren K. Hayrynen, Stefan Everling

**Author notes:** **Corresponding author** Stefan Everling, Centre for Functional and Metabolic Mapping, Robarts Research Institute, 1151 Richmond Street North, London, Ontario, N6A 5B7, Canada.

## Abstract

The common marmoset (*Callithrix jacchus*) is a small-bodied New World primate, increasing in prominence as a model animal for neuroscience research. The lissencephalic cortex of this primate species provides substantial advantages for the application of electrophysiological techniques such as high-density and laminar recordings, which have the capacity to advance our understanding of local and laminar cortical circuits and their roles in cognitive and motor functions. This is particularly the case with respect to the oculomotor system, as critical cortical areas of this network such as the frontal eye fields (FEF) and lateral intraparietal area (LIP) lie deep within sulci in macaques. Studies of cytoarchitecture and connectivity have established putative homologies between cortical oculomotor fields in marmoset and macaque, but physiological investigations of these areas, particularly in awake marmosets, have yet to be carried out. Here, we addressed this gap by probing the function of posterior parietal cortex (PPC) of the common marmoset using electrical microstimulation. We implanted two animals with 32-channel Utah arrays at the location of the putative area LIP and applied microstimulation while they viewed a video display and made untrained eye movements. Similar to previous studies in macaques, stimulation evoked fixed-vector and goal-directed saccades, staircase saccades, and eye blinks. These data demonstrate that area LIP of the marmoset plays a role in the regulation of eye movements, provide additional evidence that this area is homologous with that of the macaque, and further establish the marmoset as valuable model for neurophysiological investigations of oculomotor and cognitive control.

**New & Noteworthy:** The macaque monkey has been the preeminent model for investigations of oculomotor control, but studies of cortical areas are limited as many of these areas are buried within sulci in this species. Here we applied electrical microstimulation to the putative area LIP of the lissencephalic cortex of awake marmosets. Similar to the macaque, microstimulation evoked contralateral saccades from this area, supporting the marmoset as a valuable model for studies of oculomotor control.

## Introduction

The common marmoset (*Callithrix jacchus*) has recently gained considerable attention as a model for biomedical research in general (Mansfield, 2003) and neuroscience research in particular (Belmonte et al., 2015; Mitchell et al., 2015; Miller et al., 2016; French, 2019). Accordingly, considerable effort has been directed toward mapping the marmoset brain (Okano and Mitra, 2015), and establishing homologies between cortical areas in this species and the macaque (Paxinos et al., 2012; Solomon and Rosa, 2014; Bakola et al., 2015), which to date has been the most commonly used primate model. The marmoset model also holds substantial promise for the study of oculomotor control. Marmosets are highly visual, foveate animals that make both saccadic and smooth pursuit eye movements (Mitchell et al., 2014, 2015), and possess a lissencephalic cortex well-suited to laminar and high-density recordings (Mitchell and Leopold, 2015). In comparison to the macaque model, however, our knowledge of marmoset oculomotor areas is currently at an emergent stage, and it remains to establish the anatomical and physiological correspondences between cortical eye fields in the two species.

The cortical and subcortical circuitry controlling saccadic eye movements is perhaps the most thoroughly understood sensorimotor system in the primate brain (Munoz et al., 2000; Schiller and Tehovnik, 2001). In the macaque, these areas include the frontal eye fields (FEF) (Bruce and Goldberg, 1985) and lateral intraparietal area (LIP) (Andersen et al., 1987), both of which send direct projections to the midbrain superior colliculus (SC) (Selemon and Goldman-Rakic, 1988), an area critical for saccade generation. These cortical areas are buried deep within sulci in this species which highlights the advantages of marmosets with their smooth cortex. Here, we investigated the oculomotor properties of the marmoset posterior parietal cortex. In this species, the picture with respect to the parcellation and homology of the marmoset posterior parietal cortex (PPC) is thus far considerably less clear than is understood in the macaque. Using cyto and myeloarchitecture, Rosa et al. (2009) separated the marmoset PPC into two main subdivisions, PPd (dorsal posterior parietal) and PPv (ventral posterior parietal) (Rosa et al., 2009). Based on the similar pattern of myelination within marmoset PPd and macaque parietal cortex (Blatt et al., 1990), and the presence of large layer V pyramidal neurons, they proposed that this subregion contained the likely homologue of macaque LIP. Subsequent anatomical (Collins et al., 2005; Reser et al., 2013) and resting-state fMRI (Ghahremani et al., 2017) investigations have supported this view as they have shown triangulated connectivity of this area with the putative FEF and SC. As has been previously noted by other authors however(Solomon and Rosa, 2014), the identification of area LIP requires confirmation with electrophysiological techniques in awake animals to establish common functional properties.

Intracortical microstimulation has long been used as a tool to characterize the properties of brain areas by activating neuronal populations (see for review Tehovnik et al., 2006; Clark et al., 2011). Microstimulation of PPC in the awake behaving macaque has been shown to evoke body movements, eye blinks, and both saccadic and smooth eye movements (Fleming and Crosby, 1955; Keating et al., 1983; Shibutani et al., 1984; Kurylo and Skavenski, 1991). Thier & Andersen (1998) subsequently found that the region from which saccades could be evoked was restricted to area LIP within the lateral bank of the intraparietal sulcus, from which they observed both fixed-vector saccades, for which amplitude and direction were invariant with respect to initial eye position, and convergent or goal-directed saccades which tended to drive the eyes to a fixed goal location in space (Thier and Andersen, 1998). To date, no studies have investigated the oculomotor properties of the cytoarchitectonically defined region LIP in the common marmoset. Here, we applied for the first time electrical microstimulation to the putative area LIP within the PPC of awake behaving marmosets and monitored eye position while the animals were allowed to make unrestricted eye movements. To ensure maximal coverage of the cortical area, we implanted 32 channel Utah arrays in area LIP of two marmosets and applied stimulation trains of varying currents. Similar to previous findings in macaque monkeys, we observed eye blinks as well as both fixed-vector and convergent saccades. These data suggest that area LIP of marmoset plays a similar role to that of the macaque in the modulation of eye movements.

## Methods

### Subjects

Two male common marmoset (*Callithrix jacchus*) monkeys weighing 440 g and 451g each, and aged 2.5 and 4 years were subjects in the experiment. All experimental procedures conducted were in accordance with the Canadian Council of Animal Care policy on the care and use of laboratory animals and an ethics protocol approved by the Animal Care Committee of the University of Western Ontario. The health and welfare of animals was under the close supervision of university veterinarians.

Details of surgical procedures and training methods involved in preparation of the animals for awake behaving experiments have been previously described (Johnston et al., 2018). Briefly, each animal was acclimated to restraint within a custom-designed primate chair and subsequently underwent an aseptic surgical procedure in which a combination recording chamber/head holder was attached to the skull using dental resin and adhesive (Bisco All-Bond, Bisco Dental Products, Richmond, BC, Canada). This allowed restraint of the head within a custom-designed stereotaxic frame during experimental sessions.

Following subsequent additional training to acclimate the animals to head restraint, a second aseptic surgery was carried out in which a 32 channel Utah array (Blackrock Microsystems, Salt Lake City, UT, USA) was inserted into the putative area LIP within the posterior parietal cortex (PPC) of each monkey. Electrodes within these arrays were 1mm in length and had an inter-electrode spacing of 400 µm. During this surgery, a microdrill was used to open 4mm burr holes in the skull to allow access to PPC at locations based upon the stereotaxic coordinates of area LIP (1.4 mm anterior, 6 mm lateral) (Paxinos et al., 2012) which corresponds to the location of a posterior parietal region previously shown to have strong resting-state connectivity with the midbrain superior colliculus (SC) (Ghahremani et al., 2017). We additionally confirmed these locations visually by noting the location of a small blood vessel corresponding to the location of a shallow sulcus believed to be homologous to the intraparietal sulcus of other primate species. Arrays were manually inserted such that the width of the array straddled the sulcus and the array length covered as much of the sulcus length as possible. Following array insertion, the array wires and connector were fixed in place within the recording chamber using dental adhesive, and the array and remaining exposed cortical surface within the burr hole were covered with medical-grade silicone elastomer adhesive (World Precision Instruments, Sarasota, FLA, USA). A screw-hole was drilled into the skull posterior to the location of the implanted array to place the ground screw. The ground wire of the array was then tightly wound around the base of the screw to ensure good electrical connection. Any remaining exposed wire was then covered with additional protective layers of dental adhesive as required. After full curing of the adhesive, a removable protective cap was fixed in place on the recording chamber.

### Microstimulation Protocol

Prior to applying microstimulation, we first verified that individual sites in the microelectrode array were within cortex by monitoring for extracellular neural activity using the Open Ephys acquisition board (http://www.open-ephys.org) and digital headstages (Intan Technologies, Los Angeles, CA, USA). For both animals, spike activity began about 2 weeks following array implant surgery. Microstimulation pulses were delivered using the Intan RHS2000 Stimulation/Recording Controller system and digital stimulation/recording headstages (Intan Technologies, Los Angeles, CA, USA). Stimulation trains consisted of 0.3ms biphasic current pulses delivered at 300 Hz for a duration of 100-300ms, at current amplitudes varying between 40 and 240 µA. For those sites at which saccades were evoked, we additionally determined the threshold current, defined as the minimum current at which saccades could be evoked on 50% of stimulation applications. In sessions investigating whether saccades could be evoked at individual array sites, eye movements were uncontrolled and microstimulation trains were triggered manually by the experimenter. In those on which we investigated effects of initial eye position on saccade direction and amplitude, stimulation trains were triggered by behavior control software based on fixation location and duration criteria (see Eye Movement Recording, below).

### Eye Movement Recording

Animals were seated in a primate chair that was integrated with a custom designed stereotaxic frame for head restraint and eye movement recording. The chair/frame system was mounted on a table within a sound-attenuating chamber (Crist Instruments Co., Hagerstown, MD, USA). Their heads were restrained and a liquid spout placed at their mouth for reward delivery. Rewards consisted of acacia gum and were delivered via infusion pump (Model NE-510, New Era Pump Systems, Inc., Farmingdale, New York, USA). Eye positions were monitored via high-speed (1000 Hz) infrared video oculography which monocularly tracked pupil location (EyeLink 1000, SR Research, Ottawa, ON, Canada), and recorded together with microstimulation parameters using the Intan Simulation/Recording controller. Eye position was calibrated in each session by requiring marmosets to fixate on images of marmoset faces presented at several predetermined locations, in order to receive a liquid reward. All stimuli were presented on a CRT monitor (ViewSonic Optiquest Q115, 76 Hz non-interlaced, 1600 x 1280 resolution) at a viewing distance of 42cm. Stimulus presentation and reward delivery were carried out under computer control using the CORTEX real-time operating system (NIMH, Bethesda, MD, USA).

In the initial series of microstimulation sessions, animals freely viewed video images on the CRT monitor while we manually applied stimulation trains at single array sites and monitored the animals’ horizontal and vertical eye positions to determine whether saccades could be evoked at a given site. For those sites at which saccades could be reliably evoked, we carried out stimulation current series to determine saccade thresholds. Rewards were given manually by the experimenters to maintain the animals’ level of alertness throughout the session but were not contingent on any oculomotor behavior. To further investigate effects of initial eye position on saccade directions and amplitudes, in a subsequent series of sessions we revisited many of the sites at which saccades could be evoked and applied stimulation under computer control while the animal’s initial eye positions were at each of a set of predetermined locations. In these sessions, animals freely viewed the same video images and stimulation trains were triggered by the CORTEX real-time operating system when their eye positions were maintained within 5 × 5 degree electronic windows centred on one of a series of predetermined locations for a period of 100 ms. Locations were at an eccentricity of 7 degrees at the upper left, lower left, upper right, and lower right quadrants of the display monitor, or at the monitor centre.

### Confirming Array Location

Ex vivo MRI for marmoset B and in vivo micro-CT scan for marmoset W were conducted to confirm the positioning of the array with respect to local landmarks and the putative area LIP. As marmoset W was involved in further data experiments, a micro-CT scan was carried out to determine the array location brain of this animal.

#### Ex vivo MRI scan

Marmoset B was sacrificed at the end of the data acquisition process to prepare the brain for ex vivo MRI scan. The animal was deeply anesthetized with 20 mg/kg of ketamine plus 0.025 mg/kg Medetomidine and 5% isoflurane in 1.4-2% oxygen to reach beyond the surgical plane (i.e. no response to toe pinching or cornea touching). It was then transcardially perfused with 200 ml of phosphate buffered saline, followed by 200 ml of 10% buffered formalin. The brain was then extracted and stored in 10% buffered formalin for more than a week prior to performing the ex vivo scan. On the day of the scan, the brain was transferred to another container for imaging and immersed in a fluorine-based lubricant, Christo-lube (Lubrication Technology, Inc), to improve homogeneity and avoid susceptibility artifacts at the boundaries. The ex vivo image was then acquired using a 9.4T, 31 cm horizontal bore magnet (Varian/Agilent) and Bruker BioSpec Avance III console with the software package Paravision-6 (Bruker BioSpin) and a custom-built 15-cm-diameter gradient coil with 400 mT/m maximum gradient strength (xMR, London, Ontario, Canada; Peterson et al., 2018). An ex vivo T2-weighted image was acquired with the following scanning parameters: repetition time (TR) = 5s, echo time (TE) = 45 ms, field of view (FOV) = 40 × 32, image size = 160 × 128, slice thickness = 0.5 mm.

To identify the location of the previously implanted array, the resulting T2-weighted image was registered to the NIH marmoset brain template (Liu et al., 2018) using the registration packages of the FSL software (fMRI Software Library: http://www.fmrib.ox.ac.uk). This registration process inherently involved two steps: the NIH template was based on registration of a brain to the Paxinos et al (2012) brain; then the experimental brains were registered to the NIH template to provide the final image. Upon visual examination of the image, an indentation of comparable size to the array (2.4 mm) was identified on the surface of the cortex within the PPC that represented the array location. The location of this region of interest was interpolated on the cortical surface to create a mask across this indentation. The mask was then projected on to the surface space in CARET toolbox (Van Essen et al., 2001), using a surface-based version of the NIH volume template that was kindly provided by the authors of the NIH marmoset brain template (Liu et al., 2018). The array mask was then compared to the area LIP as defined by the parcellated regions of the NIH template, that was also projected on CARET surface space.

#### In vivo micro-CT scan

Marmoset W was imaged on a live-animal micro-CT scanner (eXplore Locus Ultra, GR Healthcare Biosciences, London, ON) to identify the location of the array on the brain. Prior to the scan, the animal was anesthetized with 15mg/kg Ketamine mixed with 0.025mg/kg Medetomidine. It was then placed on the CT bed in supine position with arms along its sides. X-ray tube potential of 120 kV and tube current of 20 mA were used for the scan, with the data acquired at 0.5º angular increment over 360º, resulting in 1000 views. The resulting CT images were then reconstructed into 3D with isotropic voxel size of 0.154 mm. Heart rate and SpO2 were monitored throughout the session. At the end of the scan, the injectable anesthetic was reversed with an IM injection of 0.025mg/kg Ceptor.

To find the location of the array with respect to the NIH template, the acquired CT image was similarly registered to the NIH marmoset brain atlas (Liu et al., 2018) using the FSL software (fMRI Software Library: http://www.fmrib.ox.ac.uk). Similar to the ex vivo MRI data, an ROI mask was created over the traces of the array across the surface of the cortex to represent the location of the array. This mask was projected on the surface space using CARET.

### Data Analysis

All eye movement analyses were carried out using custom-written codes in MATLAB software (The Mathworks, Natick, MA). Horizontal and vertical eye traces were low-pass filtered at 30 Hz and horizontal and vertical eye traces were then aligned with the stimulation onset to detect changes in eye movements following the stimulation pulse. We only considered changes in eye position that were greater than 2º and fell within a window of 200ms from the onset of microstimulation. Saccades were detected based on first order derivative, corresponding to the velocity of eye movements. The start of a saccade was defined by the first time point in which the speed of the eye movement reached 30 º/s and the first time point at which the speed went back to zero marked the end point of saccade. All saccades were normalized to the baseline by subtracting the mean of the eye position within a 50ms window before the onset of stimulation. Saccade amplitudes were defined based on the start and endpoints of saccades, as determined by the velocity criteria. The obtained values per trial were then averaged across all saccadic trials for each site of the array. Saccade latency was defined as the time from the onset of microstimulation to the onset of the saccade. The duration of saccades was calculated based on the difference between the time point of the start and end of saccades. The same procedures were employed in those sessions in which we investigated the effect of initial eye position, except that in this case saccades were not normalized to baseline (i.e. pre-saccadic eye position). To quantify the effect of initial eye position on microstimulation-evoked saccades, we applied the linear regression modeling technique proposed by Russo and Bruce (1993). For every site of interest, the size of saccades defined as the difference of saccade offset and onset, was plotted versus all different initial eye positions to produce scatter plots for horizontal and vertical saccade components, separately. A line of regression was fit into each scatter plot, the slope of which represented the effect of initial eye position on elicited saccades. A value that was closer to zero indicated that the elicited saccades had mostly unchanged trajectories, independent of the initial eye position. This is referred to as fixed-vector saccades. A value close to 1 or −1 implied that the elicited saccades changed trajectory depending on the initial eye position and converged towards a common orbital position irrespective of where the eye started. Such saccade vectors are defined as convergent or goal-directed saccades (Russo and Bruce, 1993).

## Results

### Confirming the location of the array

The results of the ex vivo MRI for marmoset B and in vivo micro-CT scan for marmoset W are demonstrated in Figure 1. The Figure on the left displays the location of the array for both marmoset B (red patch) and marmoset W (blue patch) registered on the surface space of the left hemisphere of the marmoset brain using CARET surface registration toolbox, with registration-based estimates of the cytoarchitectonic boundaries overlaid in white. The Figure on the right displays the location of the arrays in more details, by zooming into the area enclosed within the blue box as shown on the left Figure, with all the overlapping and neighboring areas labelled according to the NIH brain parcellation map (Liu et al., 2018). The purple patch in these Figures demonstrates the overlapping area between the arrays from both animals. As can be observed, the array location mostly fell within the boundaries of area LIP, while covering parts of areas MIP and VIP in both animals.

**Figure 1.**
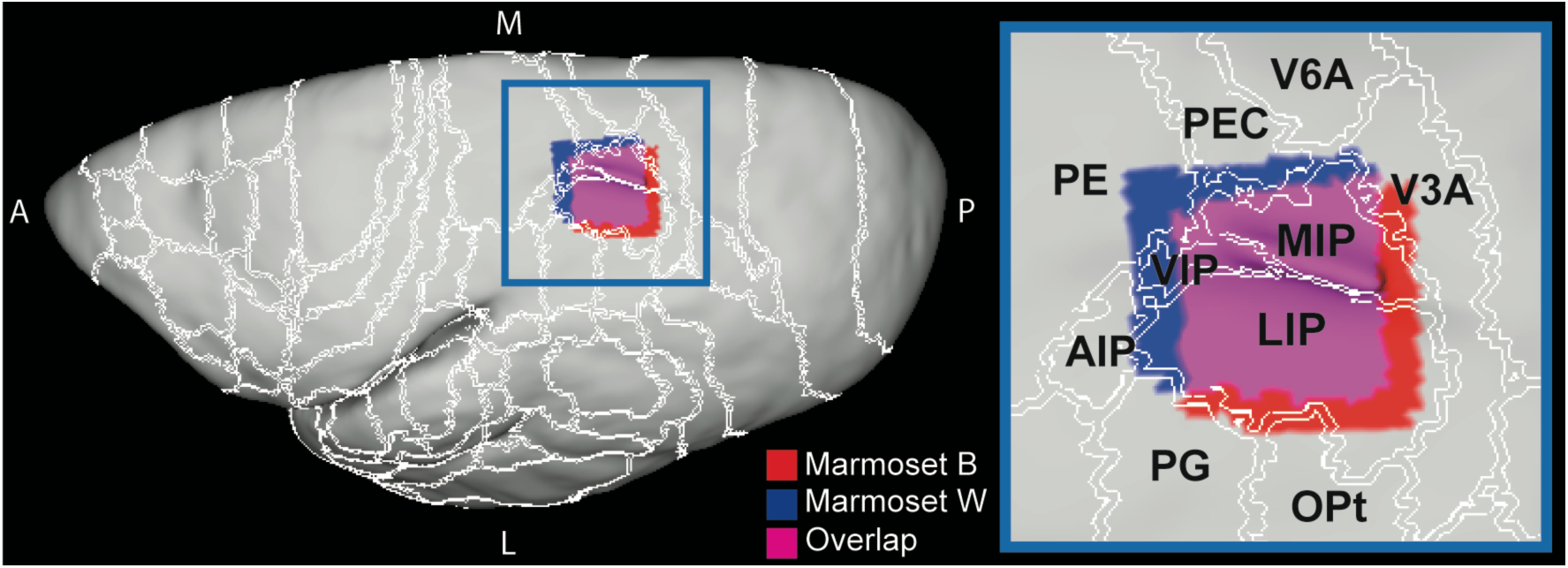
Identification of the positioning of the array using the ex vivo MRI for marmoset B and in vivo micro-CT scan for marmoset W. The Figure on the left displays the location of the array for both marmoset B (red patch) and marmoset W (blue patch) registered on the surface space of the left hemisphere of the marmoset brain, with cortical boundaries overlaid in white. The figure on the right displays the location of the arrays in more details, by zooming into the area enclosed within the blue box as shown on the left Figure, with all the overlapping and neighboring areas labelled according to the NIH brain parcellation map (Lui, 2018). The purple patch in these Figures demonstrates the overlapping area between the arrays from both animals. LIP: lateral intraparietal area, MIP: medial intraparietal area, VIP: ventral intraparietal area, AIP: anterior intraparietal area, PE: parietal area PE, PEC: caudal part of parietal area PE, PG: parietal area PG, OPt: occipito-parietal transitional area, V6A: visual area 6A, V3A: visual area 3A. M: medial, L: lateral, P: posterior, A: anterior.

### Oculomotor Effects of LIP Microstimulation

In both animals, microstimulation of putative area LIP evoked saccades and eye blinks at multiple sites of the implanted array. For Marmoset B, we observed saccades at 21/32 sites, and eye blinks at 2/32 sites. For Marmoset W, we observed saccades at 23/32 sites, and eye blinks at 9/32 sites. Figure 2A depicts this pattern of observations across all sites of the array for both animals. As can be seen from the Figure, saccades were evoked at many array sites, with the exception of the most anterior sites, from which we obtained either blinks (Marmosets W and B), or no response (Marmoset B). At some of these no response sites in Marmoset B we also never recorded any spiking activity in separate sessions (circles with cross sign in Fig. 2A). Thus, we cannot exclude the possibility that these electrodes were not in grey matter or that the cortex was damaged at those sites. The dashed line in Fig. 2 marks the approximate border line of area MIP based on the location of the shallow intraparietal sulcus (IPS) from the NIH atlas, as depicted in Fig.1. A considerable portion of the sites that elicited saccades fell on the medial side of the IPS, overlapping with the location of putative area MIP and most of the sites eliciting eye blinks overlapped with area VIP. For most sites at which microstimulation evoked saccades, we carried out current series to determine saccade thresholds, which we defined as the current at which saccades could be evoked on 50% of trials. Thresholds ranged from 40 to 240 µA. The topography of saccade thresholds is depicted in Figure 2B. We generally noted higher thresholds in Marmoset B, which was most likely attributable to the depth of the electrodes within cortex for this animal. We did not observe a clear pattern of threshold topography in either animal.

**Figure 2.**
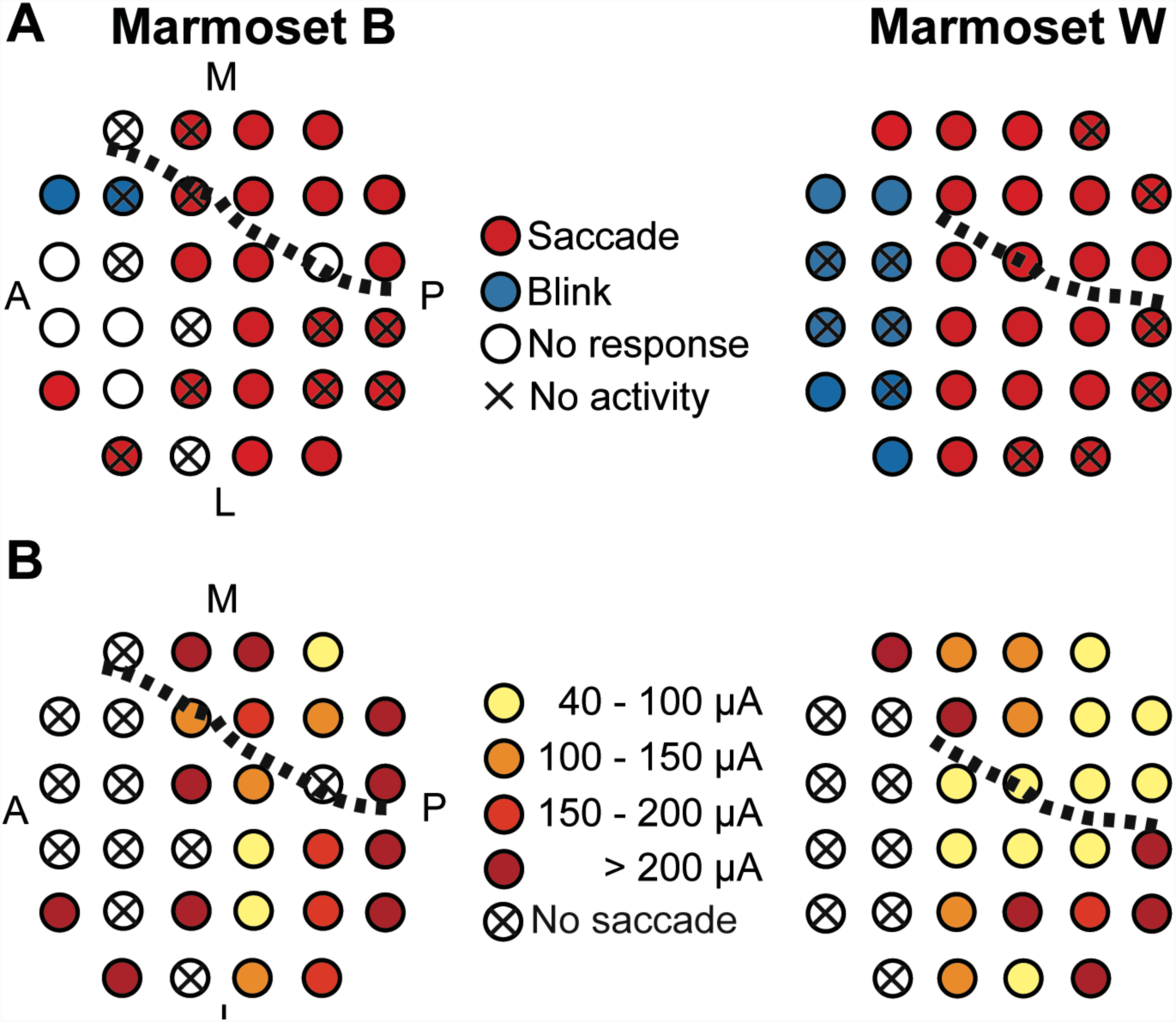
(A) Mapping the results of electrical microstimulation on individual sites of the array implanted in left area LIP for marmosets B (left) and W (right). Each circle corresponds to an electrode channel within the array. Red circles indicate sites that elicited saccades upon microstimulation. Blue circles designate sites that elicited blinks while white ones indicate sites in which no specific response was observed during microstimulation. Crossed signs refer to sites where no spiking activity was recorded. Dashed line marks the approximate border between areas LIP and MIP based on the location of the shallow intraparietal sulcus from the NIH atlas, as depicted in Figure 1. (B) Mapping the distribution of saccade onset thresholds using gradients of red. Circles with a cross sign refer to sites whose microstimulation did not evoke saccades. Dark red circles are sites with the highest saccade onset thresholds of above 200 µA. Light red marks sites with thresholds in the range of 150 to 200 µA. Orange marks sites with thresholds of 100 to 150 µA and light yellow indicates sites with the lowest saccade thresholds in the range of 40 to 100 µA. A: anterior, P: posterior, M: medial, L: lateral.

An example of a microstimulation-evoked saccade is presented in Figure 3 from a representative site in Marmoset W (marked in grey within the array grid on the top right of Fig. 3). Here, microstimulation currents of varying amplitudes (75, 100, 200 µA) elicited contralateral saccades with an upward vertical component (Fig. 3A). In general, we noted that the probability of evoking a saccade at a responsive site increased as a function of stimulation current (Fig. 3B left). Moreover, the amplitude of the evoked saccades was nearly constant across varying current amplitudes, especially past the current threshold, and fell in the range of 7.5 to 9 degrees (Fig. 3B right).

**Figure 3.**
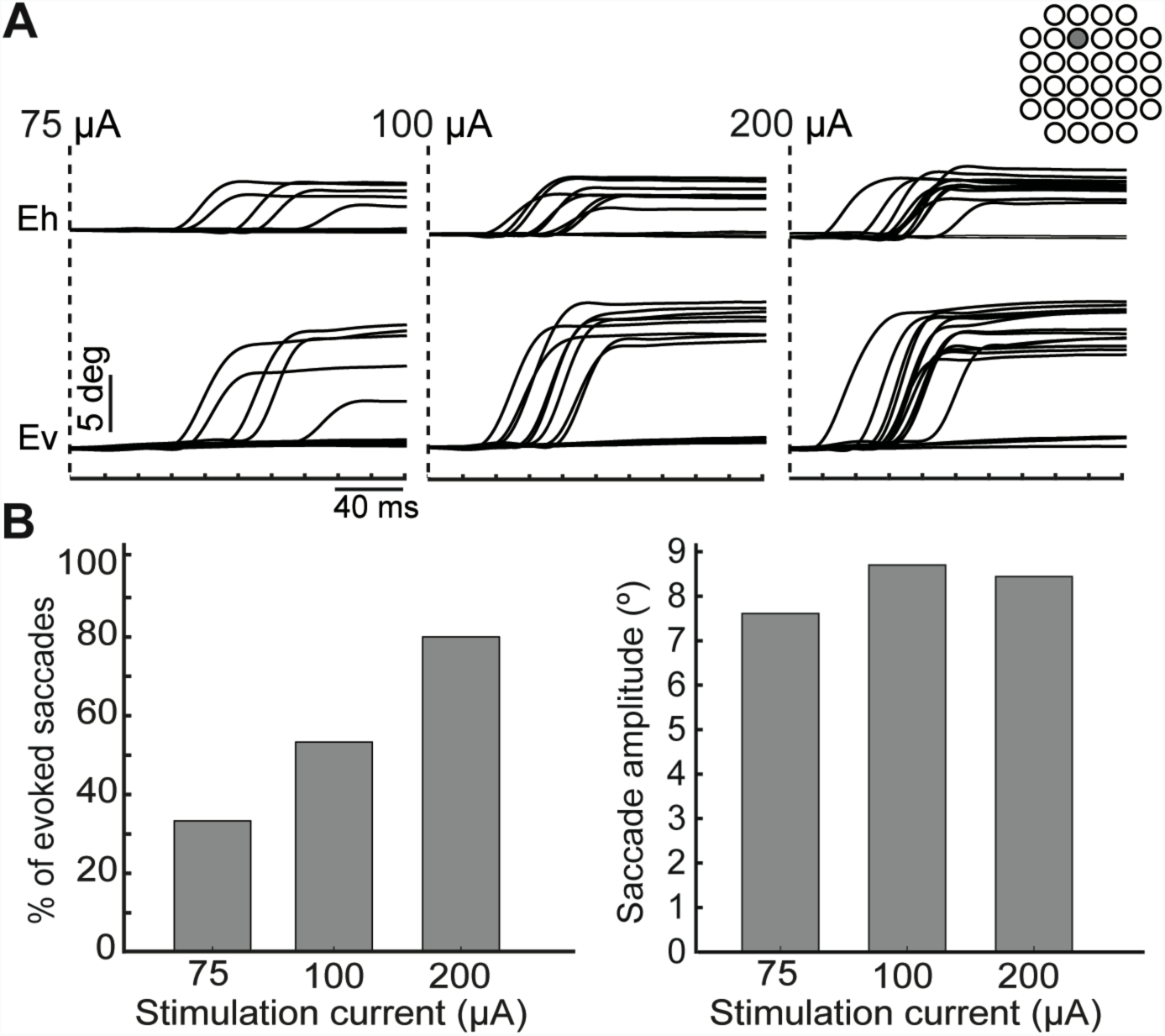
(A) Examples of saccades from a representative site (marked on the array grid on the top right) of the array in monkey W that was microstimulated at three different current amplitudes: 75 µA (left), 100 µA (middle), 200 µA (right). For each plot, the dashed line marks the onset of microstimulation and the horizonal axis represents time in milliseconds with a calibration bar that represents 40 ms. The traces at the top refer to the horizontal eye movements (E_h_), while the bottom ones refer to the vertical eye movements (E_v_). The calibration bar for the amplitude of these traces is given at 5 degrees for all plots. (B) Bar plots representing the percentage of elicited saccades (left) and their amplitude (right) at each microstimulation current amplitude (horizontal axis) from the same representative site.

### Saccade Latency and Duration

Saccade onset latency was defined as the time from the onset of the microstimulation pulse to saccade onset. Latency values were averaged across all trials of all saccade-eliciting sites for both animals and plotted as a function of the stimulation current, ranging from 50 to 240 µA (Fig. 4A). Onset latencies declined as a function of the stimulation current. Mean latencies ranged from 64 ms at 200-250 µA to 87 ms when currents were in the range of 50-100 µA. Error bars were calculated based on the standard error from the mean.

**Figure 4.**
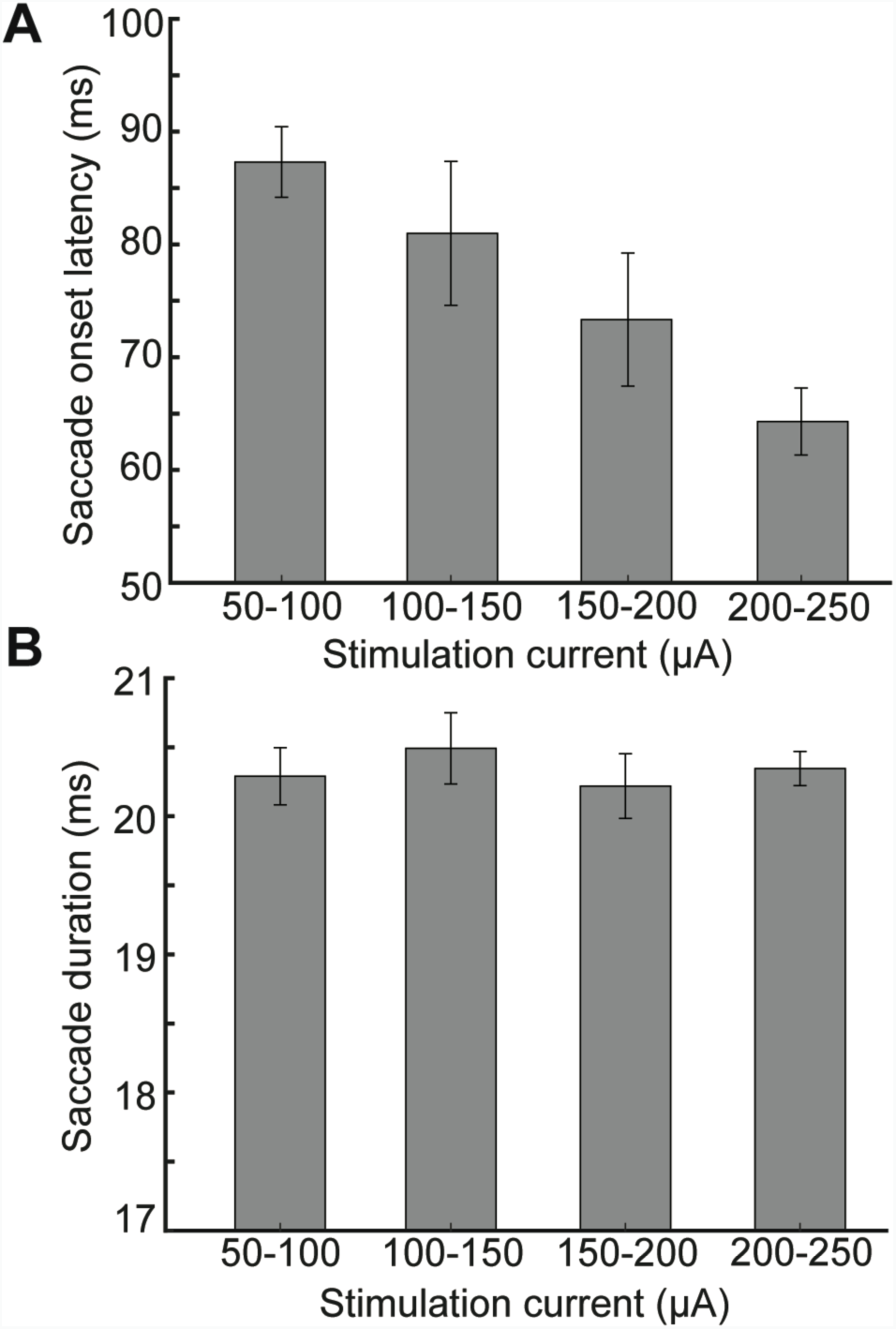
(A) Bar plot representing saccade onset latency from all saccadic sites of the array for both marmosets as a function of the amplitude of the microstimulation current. (B) Bar plot representing the duration of all saccades for both animals as a function of the amplitude of the microstimulation current. Error bars represent the standard error from the mean.

Saccade duration was defined as the time from saccade onset to the end of saccade. This parameter was similarly averaged across all trials of all saccade-eliciting sites of the array for both monkeys and plotted against stimulation current amplitude. As demonstrated in Figure 4B, saccade duration was 20 ms on average and there was no significant difference in saccade duration across different stimulation current amplitudes. Error bars were calculated based on the standard error from the mean.

### Staircase saccades

At some sites within the array, we found trials in which prolonged (300 ms) stimulation was able to evoke staircase saccades, separated by intervals of variable duration (80 to 140 ms). There were four such sites identified in marmoset W that exactly overlapped with sites from which fixed-vector saccades were elicited. There was only one site with staircase saccade identified in marmoset B. Two examples of such staircase saccades in marmoset W are shown in Figure 5 from two individual sites that were stimulated at current amplitudes of 150 and 225 µA, respectively. The amplitude of the saccades within the staircase sequence decreased in some cases (Fig. 5 left) and was less variable in other cases (Fig. 5 right), while the direction of the saccades within the staircase remained the same in most cases observed.

**Figure 5.**
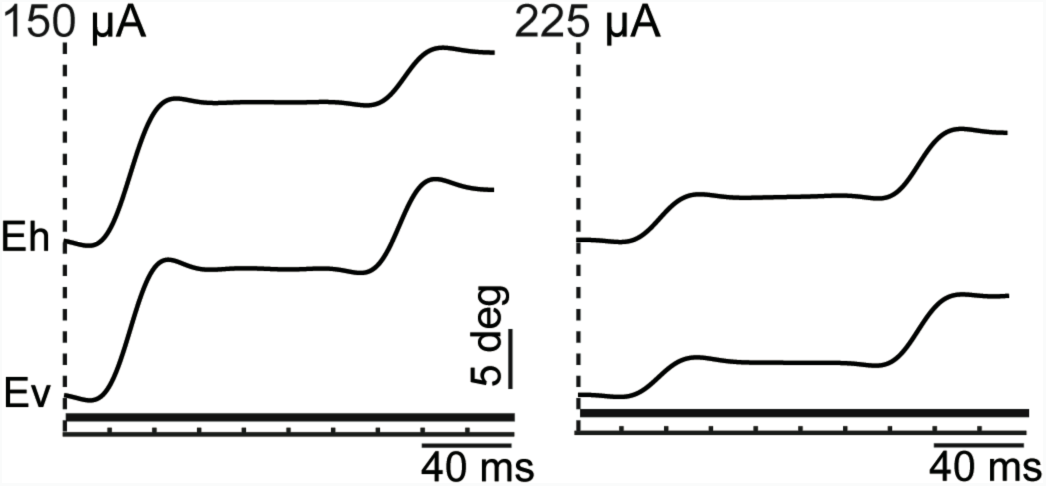
Representative examples of staircase saccades from two individual sites of the array in monkey W, stimulated at 150 µA (left) and 225 µA (right) respectively. Dashed line refers to the onset of stimulation and the solid horizontal bar shows the duration of stimulation. The horizontal axis refers to time with a calibration bar of 40 ms. The traces on the top display horizontal eye movements (E_h_) while the bottom traces show vertical eye movements (E_v_), both with an amplitude calibration bar of 5 degrees.

#### Effects of initial eye position

Based on the previous literature on macaque LIP, it has been reported that one of the factors influencing the direction or amplitude of saccades induced by LIP microstimulation is the initial position of the eye at the time of microstimulation (Shibutani et al., 1984; Kurylo and Skavenski, 1991; Thier & Andersen, 1996; Their & Andersen, 1998). To investigate this in the marmoset PPC, in a subset of sessions and sites we applied microstimulation whenever the animal’s eye position fell within one of 7 predefined zones. Next, we applied regression analysis to each of these sites for horizontal and vertical saccade components individually, to examine the impact of initial eye position on the convergence of the elicited saccades. The slope of the regression line was calculated and used as an index to indicate the level of the impact (Russo and Bruce, 1993). Out of the 10 saccadic sites investigated, three sites exhibited saccades for which amplitude and direction were modified such that eye position was driven to converge on a common location, from any starting position across the display. An example of such sites is shown in Figure 6A (right) from a representative site of the array in marmoset W with its corresponding regression analysis plots (6B, right). The location of these sites is marked within the array grid as displayed on the top right of each plot. The preferred “goal zone” was generally located in the contralateral upper visual field as demonstrated in the representative example of Figure 6A (right). The slope of the regression line for horizontal saccade component (K_h_, blue) in this site was 0.24 and for vertical component (K_v_, red) was 0.65. Most of the remaining sites exhibited fixed-vector saccades that maintained similar amplitudes and directions irrespective of the starting position of the eye at the onset of microstimulation. A representative site that elicited such fixed-vector saccades in marmoset W is shown in Figure 6A (left) with its corresponding regression analysis plot (6B, left). Here, the slope of the regression line for horizontal saccade component (K_h_, blue) in this site was 0.02 and for vertical component (K_v_, red) was −0.03, which were much closer to 0 compared to the representative site for the goal-directed saccade (Fig. 6A, right). There were few sites that exhibited saccades that did not exactly belong to either category. However, all evoked saccades (vector or goal-directed) exhibited an upward bias, contralateral to the site of stimulation. Figure 7 displays the slopes of the regression line for horizontal (left, *K*_*h*_) and vertical (right, *K*_*v*_) saccade components plotted as bar graphs for all the 10 sites in which the effect of initial eye position was investigated. The location of these sites is marked on the array grid as shown on the top right of Figure 7. For the three sites that seemed to elicit goal-directed saccades, *K*_*h*_ was in the range of 0.14 to 0.25, which was comparable to other sites (Fig. 7A). However, in these three sites *K*_*v*_ varied from 0.5 to 0.6 which was considerably higher than the slope of the vertical saccade components of all the other sites (Fig. 7B). In the case of remaining sites, the slope of the regression line for horizontal saccade components (*K*_*h*_) ranged from 0.01 to 0.2 and for vertical saccade components (*K*_*v*_) ranged from −0.1 to 0.37.

**Figure 6.**
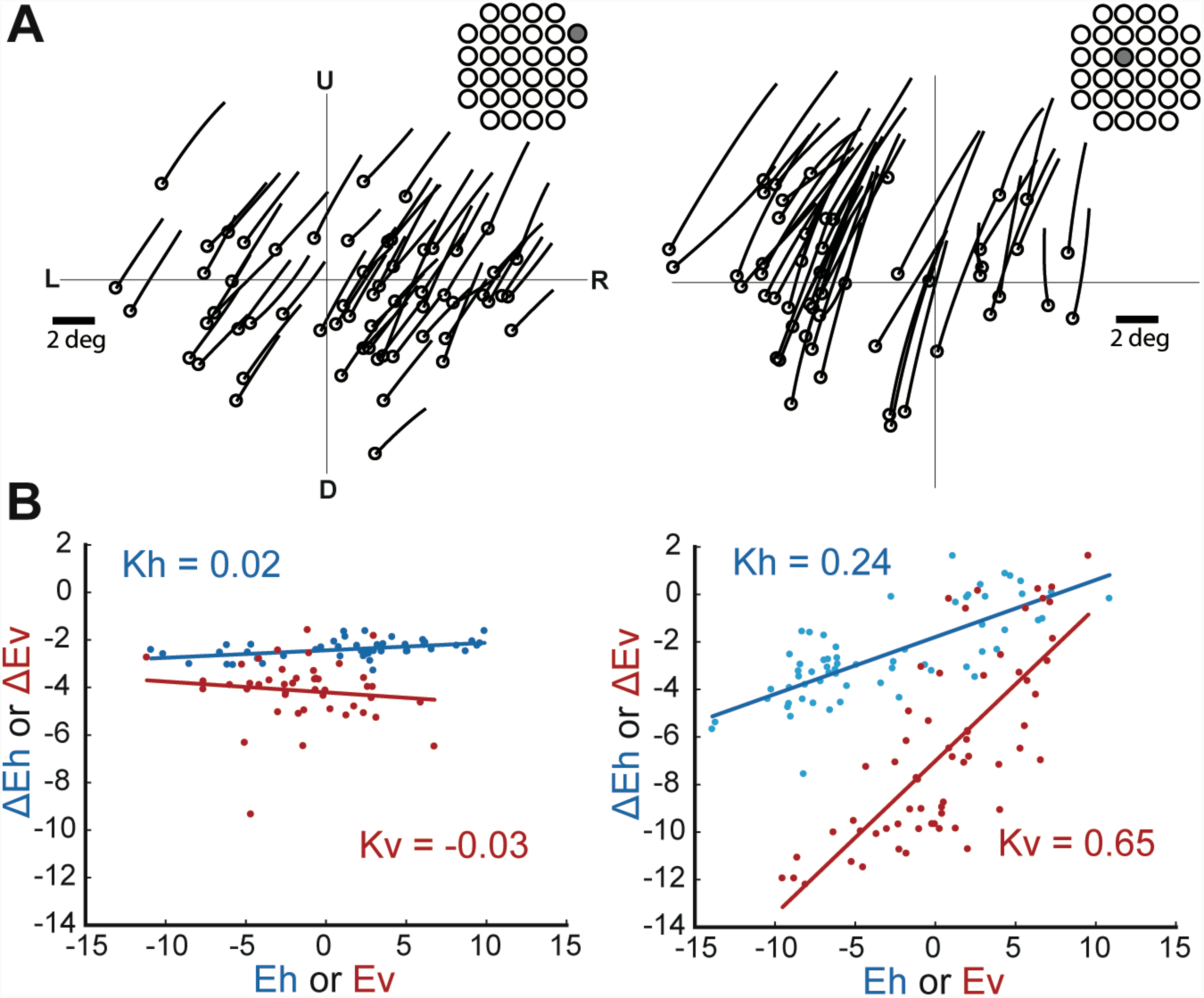
Effect of initial eye position on stimulation-evoked saccades. (A) Examples of the effect of initial eye position on evoked saccades from 2 representative sites of the array in monkey W as marked on the array grid. Left: fixed-vector saccades, right: goal-directed saccades. Circles indicate the starting position of saccades. Vertical and horizontal axes illustrate the distribution of starting position of saccades across the display. L: left, R: right, U: up, D: down. Saccade amplitude calibration bar is given for 2 degrees. (B) Corresponding regression plots for the representative sites displayed on part A. Each Figure displays two scatter plots for each of the horizontal (K_h_, blue) and vertical (K_v_, red) saccade components, with the vertical axis being the size of saccades defined as the difference of saccade offset and onset (ΔE_h_: horizontal, ΔE_v_: vertical), versus different initial eye positions (E_h_: horizontal, E_v_: vertical). Dots represent saccade trials from that representative site and the line represents the regression line of best fit.

**Figure 7.**
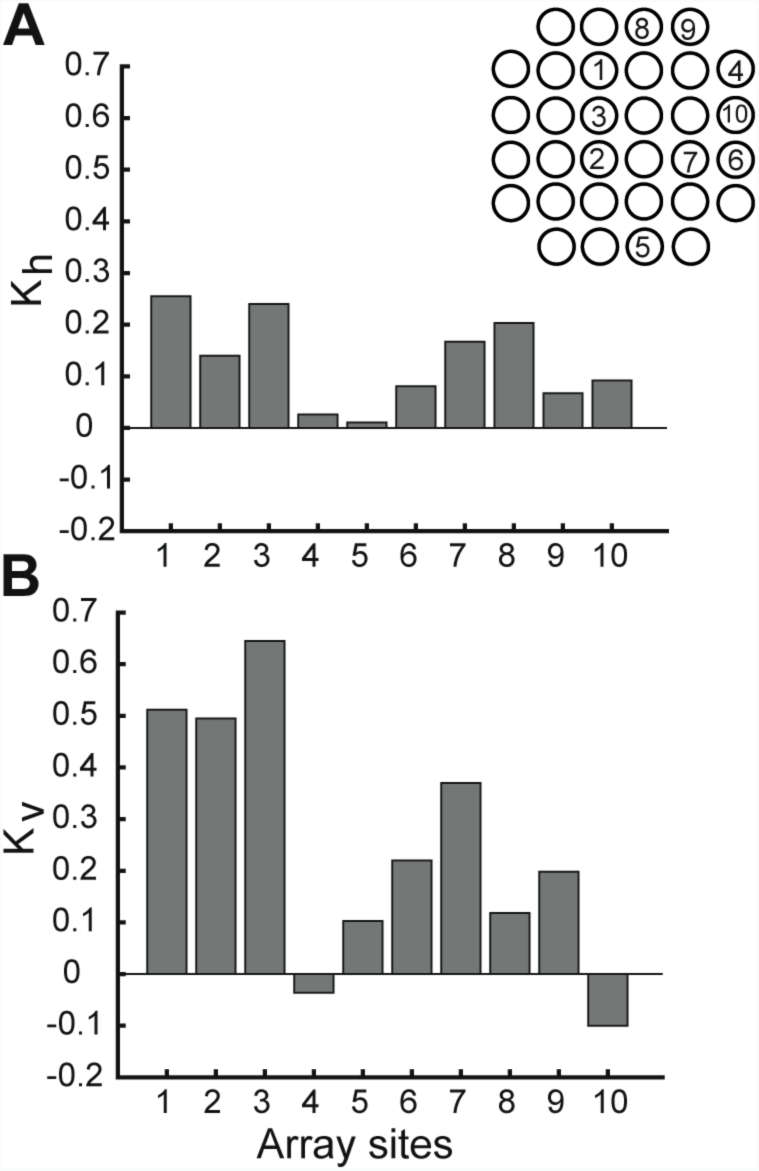
Bar plot representing the results of regression analysis from all the 10 sites of the array in which the effect of initial eye position was examined in monkey W. These specific sites are marked on the array grid as shown on the top right of the Figure. (A) bar plots representing slopes of horizontal saccade components (K_h_),(B) bar plots representing slopes of vertical saccade components ((K_v_). Horizontal axis represents individual sites and vertical axis represents slopes (left: horizontal, right: vertical).

### Saccade Direction and Amplitude

Saccade direction and amplitude were calculated based on the slope and amplitude of horizontal and vertical eye traces, between the start point and end point of saccades for each saccadic site of the array. These parameters were represented as arrows with their associated direction and amplitude. To determine the general topography of saccade directions and amplitudes, all saccades evoked at each specific site were averaged together, resulting in an average saccade vector for that site. This procedure was repeated for all saccadic sites of the array for each animal and the resulting saccade vectors were mapped onto their associated location within the array (Figure 8A). The result demonstrated that the majority of microstimulation-evoked saccades exhibited a prominent upward bias, contralateral to the site of stimulation, in both animals across most sites of the array (Figure 8A). There were few sites in Marmoset B that exhibited downward saccades. However, the horizontal component of the evoked saccades was always contralateral to the site of stimulation. In terms of the saccade amplitude, varying amplitudes were observed across array sites in both animals with more centrally located sites having larger amplitudes in general.

**Figure 8.**
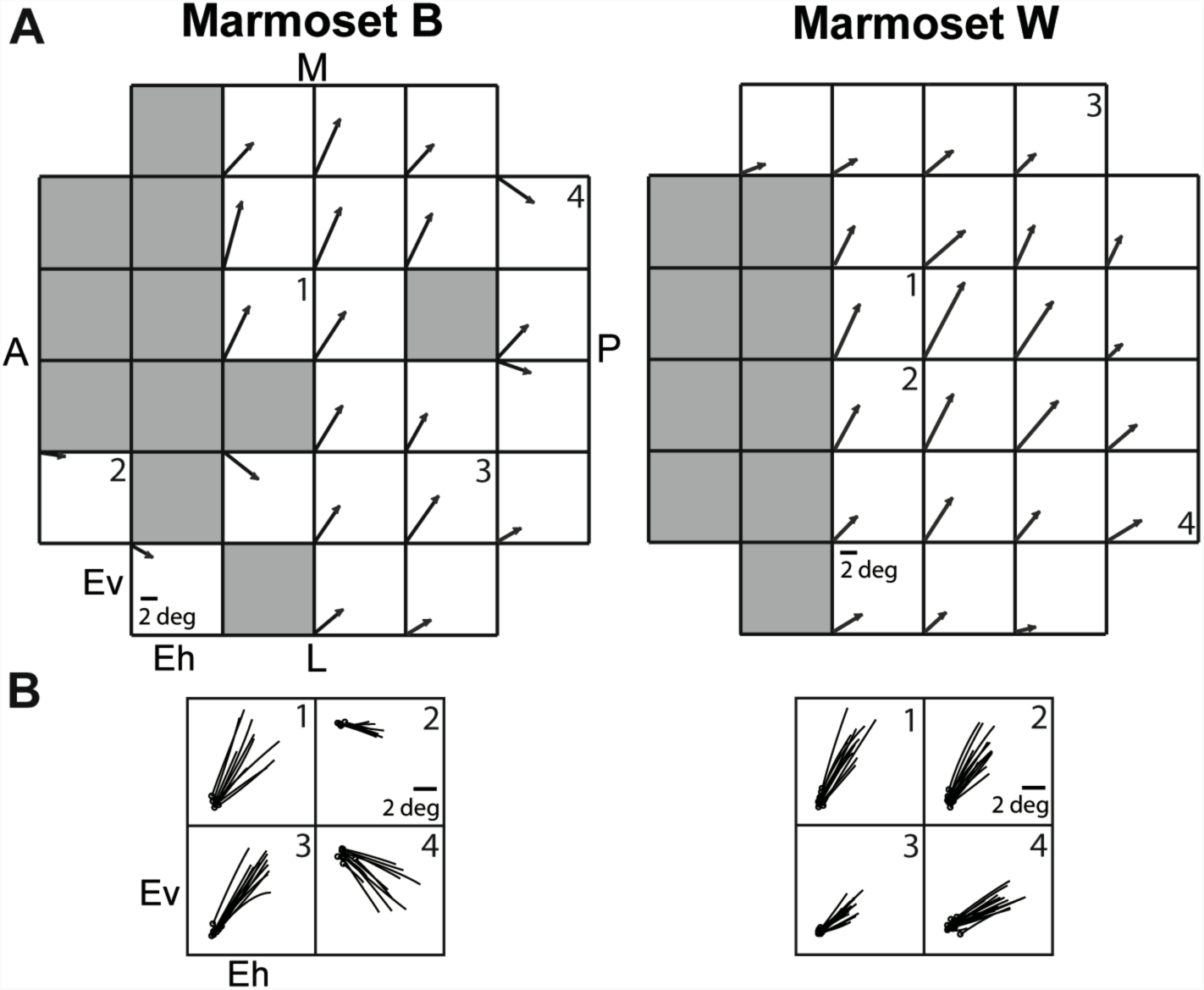
(A) Mapping the average saccade vectors across all saccadic sites of the array for marmosets B (left) and W (right). Each square corresponds to an array electrode. Grey squares refer to sites that did not elicit saccades upon microstimulation. Each arrow represents the average saccade vector evoked across all saccade trials, with its associated amplitude and direction. The amplitude of the saccades is measured by the calibration bar given for 2 degrees. Each arrow is plotted based on vertical and horizontal axes that represent vertical (E_v_) and horizonal (E_h_) eye movements, respectively. (B) Examples of individual saccades constituting an average saccade vector from four representative array sites per animal, marked by numbers 1, 2, 3, and 4 in (A). Saccade amplitude calibration bar is given for 2 degrees. Circles indicate the starting position of saccades. For each site, vertical and horizontal axes refer to vertical (E_v_) and horizontal (E_h_) eye movements, respectively. A: anterior, P: posterior, M: medial, L: lateral.

## Discussion

Here, we applied intracortical microstimulation to a subregion of the PPC, putative area LIP, to investigate the oculomotor properties of this area in common marmosets. We observed a suite of oculomotor responses including fixed-vector saccades, goal-directed craniocentric saccades, and staircases of saccades. In all cases, saccades were directed toward the hemifield contralateral to the site of stimulation, and predominantly toward the upper visual field. In some cases, we also observed eye blinks. These findings are consistent with previous electrical stimulation studies of PPC in the macaque model (Shibutani et al., 1984; Kurylo and Skavenski, 1991; Thier and Andersen, 1996, 1998), and support the view that LIP of the marmoset has similar oculomotor properties as that of macaque.

A consistent finding of microstimulation experiments in macaque LIP is the presence of two broad classes of saccades: so-called “modified vector” or retinotopic saccades with relatively consistent directions and amplitudes regardless of the initial position of the eyes, and “goal-directed” or craniocentric saccades, the amplitudes and directions of which vary with initial eye position such that they tend to be driven toward a particular goal location (Kurylo and Skavenski, 1991; Thier and Andersen, 1996, 1998). Sites at which these saccade types can be evoked have a topographic distribution, such that vector saccades are evoked at more caudal sites, while goal-directed saccades are confined to a small rostral region in the floor of the of the intraparietal sulcus and extending into the medial bank, termed the “intercalated zone” (Thier and Andersen, 1996, 1998). We similarly observed both classes of saccades following microstimulation of marmoset putative area LIP. We found, however that goal-directed saccades were rare and not restricted to a specific cluster of sites. One possible explanation for the relative lack of goal-directed sites we observed is the limited oculomotor range of the marmoset. In some cases, the goal-directed saccades evoked from macaque intraparietal sulcus drove the eyes to a goal location beyond the range of ocular motility (Kurylo and Skavenski, 1991; Thier and Andersen, 1996). In marmosets, very few saccades occur beyond about 12 degrees during head-fixed free viewing (Mitchell et al., 2014; Mitchell and Leopold, 2015), which is considerably less than the approximately 30 degree eccentric range of the macaque (Tomlinson and Bahra, 1986; Heiney and Blazquez, 2011). Marmosets also rely more on movements of the head to shift gaze (Mitchell et al., 2014). It thus seems possible that we failed to observe convergent eye movements due to the fact that we were able to analyse only the initial few degrees of the eye trajectories of gaze shifts converging well outside the oculomotor range, and thus underestimated the number of sites at which goal-directed saccades could be evoked. Alternatively, such sites may simply be more rare and widely distributed in marmoset PPC. Consistent with this idea, we found that eye blinks but not goal-directed saccades could be evoked at the most rostral sites in both animals. In contrast, in macaque eye blinks and goal-directed saccades could both be evoked from the intercalated zone in the rostral portion of area LIP (Thier and Andersen, 1996, 1998). Whether this difference in co-localization of responses represents a real phylogenetic difference in modularity of PPC organization (Krubitzer et al., 1995) between these primate species remains to be definitively determined, though we noted also a continuity of sites from which saccades could be evoked in marmosets, contrasting with the typical “fractured” distribution of sites in macaque LIP, in which sites from which saccades can be evoked are organized in clusters separated by non-responsive bands of cortex (Thier and Andersen, 1996).

In contrast to differences in the distribution of goal-directed saccades, we observed a similar topography of saccade directions in marmoset area LIP to previous findings in macaques (Thier and Andersen, 1998). The directions of evoked saccades were toward the contralateral hemifield in all cases in both animals, and in the vast majority of cases these saccades had an upward component. A small number of sites with a downward component were observed in marmoset B, either at the most rostral or caudal sites. Amplitudes of evoked saccades varied from 3 to 12 degrees. In neither animal did we observe a clear topographic organization of either of these saccade parameters. In macaque LIP, microstimulation has been shown to evoke contralateral saccades with a strong upward bias and no clear organization with respect to direction or amplitude (Kurylo and Skavenski, 1991; Thier and Andersen, 1996, 1998). One potential explanation for how rarely downward saccades were evoked is that these types of saccades may be encoded in a different area within the marmoset PPC that was not covered by our implanted array. Another possibility is that many of small marmosets’ predators like raptors attack from above (Oliveira and Dietz, 2011) and therefore there might be a stronger and faster upper visual field representation in these primates ’oculomotor regions. Such over-representation of the upper visual field has been previously reported in rodents (Drager and Hubel, 1976) and more recently in primates (Hafed and Chen, 2016). Hafed and Chen (2016) reported sharper, stronger, and faster upper visual field representation in the SC of macaque monkeys. They hypothesized that SC organization is in tune with environmental constraints on eye movement exploration between the upper and lower visual fields. This was evident by a significant asymmetry observed across the horizontal meridian, such that the SC generated more accurate and lower latency saccades towards the upper visual fields (Hafed and Chen, 2016). Another study characterizing response fields of FEF neurons reported that about 70% of them were upper visual field neurons (Mayo et al., 2015). These findings suggest that the over-representation of the upper visual field may be common in other areas involved in saccade and visual exploration such as area LIP in primates.

Previous studies investigating the effects of intracortical microstimulation on motor responses in marmosets have reported that the thresholds for evoking movements is greater in this species than macaques (Burman et al., 2008), possibly due to the smaller soma size and hence decreased electrical excitability of pyramidal neurons in marmosets (Nudo et al., 1995). We systematically obtained thresholds at most PPC sites from which we were able to evoke eye movements and found that thresholds varied from 40 to 240 μA, values similar to those observed in previous studies of macaque PPC which in the literature have typically ranged from 25-200 μA (Shibutani et al., 1984; Kurylo and Skavenski, 1991; Thier and Andersen, 1998). This similarity is perhaps surprising as the Utah arrays we used here did not allow us to optimize the cortical depths at which stimulation was applied by moving individual electrodes. However, the higher current threshold seems to be a characteristic feature of marmoset area LIP compared to other oculomotor frontal areas. In fact, in a recent microstimulation study by Selvanayagam et al (2019) from our group, 1mm-length Utah arrays were implanted in frontal cortical areas of marmosets and microstimulation within the putative area FEF could elicit saccades at current thresholds as low as 12 μA. Thus, although the fixed length of Utah electrode arrays did not allow optimal targeting of layer V output neurons in either study, marmoset area LIP seems to have a higher saccade thresholds than marmoset FEF (Selvanayagam et al., 2019). We also observed that the amplitude of evoked saccades did not vary greatly across varying amplitudes of the stimulation current, especially above the threshold current. Similar findings were reported in microstimulation studies of area LIP (Shibutani et al., 1984; Kurylo and Skavenski, 1991), as well as the SC (Schiller and Stryker, 1972), and FEF (Robinson and Fuchs, 1969) in macaques. We did note however that the latencies of evoked saccades were greater following stimulation of marmoset LIP than most prior studies in macaque. We obtained latencies of 64-87ms, in comparison to the mean latencies of 30 and 50 ms found by Thier & Andersen (1998), and Shibutani et al. (1984), respectively. Whether this represents a true species difference in the role of area LIP in the control of saccades or is simply a reflection of reduced soma and axon sizes in the smaller New World marmoset remains to be determined. Studies investigating the response properties of single LIP neurons during saccades, as well as studies in the marmoset FEF should prove illuminating in this regard. Saccade duration in the present study was around 20 ms which greatly overlaps with the duration of spontaneous saccades previously reported for common marmosets (Mitchell et al., 2014), though it most closely resembles faster spontaneous saccades. Kurylo and Skavenski (1991) obtained similar findings in macaque monkeys, where there was substantial overlap in the duration of spontaneous and electrically induced saccades. They claimed that the larger spread in the duration of spontaneous saccades compared to stimulation-evoked saccades can explain the small differences in duration.

Many previous studies applying microstimulation to macaque PPC have observed evoked movements of not only the eyes, but also the pinnae, face, arms, and shoulders (Fleming and Crosby, 1955; Shibutani et al., 1984; Kurylo and Skavenski, 1991; Thier and Andersen, 1998; Cooke et al., 2003). Although we monitored the animals here for these effects, we did not observe any such movements following stimulation, even at higher current amplitudes. This may be at least partially explained by the size and locations of the implanted microelectrode arrays. Each array was implanted on the basis of resting-state fMRI and the local sulcal landmark such that it was placed roughly straddling the sulcus in mediolateral extent and centred with respect to the sulcus in rostrocaudal extent, in order to cover as much of the putative area LIP as possible. In marmoset, area LIP extends approximately 3.5 – 4mm rostro-caudally, and 2 mm mediolaterally. The dimensions of our arrays were 2.4 × 2.4 mm, and seemed to be rostrocaudally located predominantly within the borders of area LIP, and mediolaterally overlapping with area MIP on the medial side of the intraparietal sulcus (Fig. 1). Thier and Andersen (1998), found that within macaque area LIP itself, movements other than those of the eyes were evoked only in the intercalated zone from which goal-directed saccades could be evoked, at the most rostral extent of LIP. While the medial side of the IPS (area MIP) in macaques have been implicated to specifically hold a representation of hand or reaching movements towards a visual target, there is a small percentage of neurons within macaque MIP that also respond in relation to saccadic eye movements (Snyder et al., 1997, 2000). In the present study, even though the implanted array covered parts of area MIP according to the Paxinos et al. (2012), no movements other than those of the eyes were evoked at the more caudal sites from which fixed-vector saccades were evoked. Since we found that fixed-vector saccades were evoked at the majority of sites in both animals, it seems reasonable to suggest that most of our stimulation sites were restricted to the region specific to fixed-vector eye movements, and simply did not reach cortex from which non-eye movements could be evoked. This observed discrepancy between marmosets and macaques may be due to the variations of borders across individuals and the imprecisions inherent to the registration process. It can also imply that the region designated as the putative area MIP in marmosets according to Paxinos et al. (2012), is actually an extension of area LIP, since it does not seem to serve a different function compared to the LIP. In macaque monkeys, area LIP is the only parietal area with direct projections to the SC (Lynch et al., 1985; Andersen et al., 1990) and there are reciprocal connections between the FEF and the LIP and VIP, but not MIP (Stanton et al., 1995, 2005). However, previous tracing and functional connectivity studies in marmosets have demonstrated that all three intraparietal areas LIP, VIP and MIP are reciprocally connected with the putative marmoset FEF (Reser et al., 2013; Majka et al., 2016; Ghahremani et al., 2017). Accordingly, corticotectal neurons in marmoset PPC seem to be more distributed compared to the macaque (Collins et al., 2005). Although tracer studies supporting the identity of marmoset area LIP have been based on injections placed in other areas (Rosa et al., 2009) and no study involving direct injections into LIP has been published yet, open access data provided by Majka et al (2016) (marmosetbrain.org) confirm that the putative area LIP in marmosets is indeed mostly connected to visual and frontal oculomotor areas such as the FEF. Therefore, a macaque LIP homologue certainly exists within the marmoset PPC, but its precise boundaries remain unclear, such that the physiological area LIP as defined by Paxinos et al (2012) may be larger than what was originally proposed. On the other hand, we did observe eye blinks at a few stimulation sites in both animals that possibly covered parts of area VIP, consistent with observations in area VIP in macaques, a region from which defensive movements of the arms and face can also be evoked (Cooke et al., 2003). Future work using larger arrays covering a greater extent of PPC or greater cortical sampling by other means would definitely address this discrepancy in findings.

The recent increase in popularity of the marmoset model has been paralleled by anatomical studies aiming to establish homologies between cortical areas of the marmoset and rhesus macaques, which historically have been the most commonly used primate model in neuroscience research (Solomon and Rosa, 2014). In a similar vein, behavioural work has demonstrated that marmosets display many similarities with macaques with respect to their visual and oculomotor behaviour (Mitchell et al., 2014; Johnston et al., 2018, 2019). To date however, few studies have investigated the properties of cortical areas involved in oculomotor control in awake behaving marmosets, which is a critical step in determining the function of these areas and relating them to macaques and ultimately human brain function. It has been proposed that homology between cortical fields can be established on the basis of three primary criteria: cytoarchitecture, connectivity, and neural response properties (Kaas, 1987; Krubitzer, 1995). With respect to area LIP, corresponding aspects of cytoarchitecture between macaques and marmosets such as dense myelination and the presence of large layer V pyramidal neurons have been established (Bock et al., 2009; Rosa et al., 2009). Similarly, area LIP shares a common pattern of connectivity across species. This area was originally defined in macaques as a subregion within the intraparietal sulcus with extensive interconnections to the FEF and SC (Andersen et al., 1985). Such connectivity has been established in marmoset on the basis of anatomical (Collins et al., 2005; Reser et al., 2013) and resting-state fMRI (Ghahremani et al., 2017). Our data here suggest that the functional properties of LIP in both species are also similar. Although we noted some differences in the proportions and distributions of the two types of microstimulation-evoked saccades in the two species, we found that saccades could be evoked at similar currents, at marginally longer latencies, and with a similar direction bias toward the upper contralateral visual field as previous studies in macaques. Although single neuron recordings in area LIP of marmosets trained to perform oculomotor tasks are needed to definitively establish correspondence in neuronal response properties, taken together, these three existing lines of convergent evidence provide compelling support for the notion that marmoset LIP is homologous with that of macaque monkeys, and establish further the marmoset as a promising new model for the study of oculomotor and cognitive control.

## Author Contributions

MG performed experiments, analysed data, prepared the Figures, and wrote the first draft of the manuscript. KJ designed experiments, performed experiments, and edited the manuscript. LM helped with the surgery, performed perfusion and brain extraction and edited the manuscript, LS performed experiments and edited the manuscript. SE designed experiments, performed surgeries, edited the manuscript, and approved the final version of the manuscript.

## Acknowledgements

We thank N. Hague, C. Vander Tuin, W. Froese, and K. Faubert for surgical assistance and care of the experimental animals. This research was supported by the Canadian Institutes of Health Research grant FRN148365 to S.E. and the Canada First Research Excellence Fund to BrainsCAN.

